# “Gap hunting” to characterize clustered probe signals in Illumina methylation array data

**DOI:** 10.1101/059659

**Authors:** Shan V. Andrews, Christine Ladd-Acosta, Andrew P. Feinberg, Kasper D. Hansen, M. Daniele Fallin

## Abstract

**Background:** The Illumina 450K array has been widely used in epigenetic association studies. Current quality-control (QC) pipelines typically remove certain sets of probes, such as those containing a SNP or with multiple mapping locations. An additional set of potentially problematic probes are those with DNA methylation (DNAm) distributions characterized by two or more distinct clusters separated by gaps. Data-driven identification of such probes may offer additional insights for downstream analyses.

**Results:** We developed a procedure, termed “gap hunting”, to identify probes showing clustered distributions. Among 590 peripheral blood samples from the Study to Explore Early Development, we identified 11,007 “gap probes”. The vast majority (9,199) are likely attributed to an underlying SNP(s) or other variant in the probe, although SNP-affected probes exist that do not produce a gap signals. Specific factors predict which SNPs lead to gap signals, including type of nucleotide change, probe type, DNA strand, and overall methylation state. These expected effects are demonstrated in paired genotype and 450k data on the same samples. Gap probes can also serve as a surrogate for the local genetic sequence on a haplotype scale and can be used to adjust for population stratification.

**Conclusions:** The characteristics of gap probes reflect potentially informative biology. QC pipelines may benefit from an efficient data-driven approach that “flags” gap probes, rather than filtering such probes, followed by careful interpretation of downstream association analyses. Our results should translate directly to the recently released Illumina 850K EPIC array given the similar chemistry and content design.

## Background

DNA methylation (DNAm) is a type of epigenetic mark and term commonly used to denote the covalent addition of a methyl or hydroxymethyl group to a cytosine nucleotide base in the DNA sequence, typically at cytosine-guanine dinucleotide sequences, or CpG sites. DNAm is a necessary component to cellular differentiation during development, and is a leading mechanism for the plasticity of the genome in response to various environmental stimuli during the life course [1]. There is an ever-increasing focus on various studies of DNAm, which can be broadly classified into three main domains: those seeking to discover the relationship between DNAm and various adverse health outcomes [2–4], those seeking to find DNAm changes associated with environmental exposures [5–7], and those screening for genetic loci that control states of DNAm (methylation quantitative trait loci, meQTLs) [3, 8]. These three groups of studies constitute the now burgeoning field of epigenetic epidemiology.

The Illumina HumanMethylation450 BeadChip (450k) has largely enabled the fast growth of epigenetic epidemiology because it effectively balances sample throughput and cost with epigenome coverage. Specifically, the 450k allows for the efficient interrogation of roughly 485,000 CpG sites in the human genome, covering 99% of RefSeq genes, CpG islands, lower-density CpG regions, termed shores and shelves, shown to be associated with differentiation and disease [9, 10], and other high value content such as microRNA promoter regions and DNase hypersensitive sites [11]. Probes are characterized by 3 distinct features: a CpG site of interest, a single-base extension (SBE) that incorporates a fluorescently labeled nucleotide for detection, and an additional 48 or 49 base pairs. The chemistry involves two probe types. Type I uses two probes per interrogated CpG site, one for a methylated sequence and one for unmethylated sequence, with measurement based on signal from a single color channel (red or green) determined by the nucleotide base incorporated via SBE. Type II probes use a single probe with measurement based on the ratio of red and green signal intensities (a two-color array rather than one-color) [11]. In this design the C base of the CpG site overlaps with the SBE site.

As use of the 450k has become increasingly widespread, there have been several contributions that have increased our general understanding of probe behavior on the 450k. One frequently cited example is that of ambiguously mapping probes, or probes that can hybridize to multiple places in the genome. A list of these probes has been made publicly available, and they are often removed prior to association analysis [12]. Several studies have also noted the existence of probes in which genetic polymorphisms may be present at the target CpG site, at the SBE, and/or elsewhere in the probe [13, 14]. Estimates of the proportion of polymorphic CpG sites out of all those interrogated by the 450k Array have ranged from 4.3% [13] to 13.8% [12]. Typically, 450k Array-based studies account for the presence of polymorphisms by using various reference annotation schemes; examples include those developed from the Database for Single Nucleotide Polymorphisms (dbSNP) [3], from the 1000 Genomes Project [8], or from the Illumina-provided manifest [15]. A recent report recommended removal of 190,672 probes (39% of the 450k Array) prior to association analysis [16] based on concordance between whole genome bisulfite sequencing data and 450k data in several potentially problematic groups of probes compared to a “high quality” group, each defined via reference annotation. However screening for potentially problematic probes based solely on pre-defined reference annotation tables can be problematic because they can vary according to the database chosen (dbSNP, 1000 Genomes, etc.), contain very rare variants, or may not be relevant to the population being investigated in a particular study. These factors could result in the misclassification of probes as being polymorphism-affected or not, and suggest against the blind removal of problematic probes classified in any part by the reference annotation method.

Recently, Daca-Roszak et al. overcame these reference annotation limitations on a small scale through the analysis of combined study-specific genotype and 450K array data on 96 probes that distinguished European and Chinese populations. 69% of these probes contained study-specific SNPs that were ancestry informative. They specifically note the existence of tri-and bi-modal beta value distributions at many of these 96 probes, and carefully delimit, through consideration of bisulfite conversion and probe chemistry, how each possible SNP at the C and G sites of interest (C/T, C/G, C/A or G/T, G/C, G/A) can affect methylated and unmethylated signal and the subsequent beta value calculated. Ultimately, the authors recommend a careful consideration of the potential influence of genetic polymorphism on DNAm signal when interpreting epigenome-wide association study (EWAS) results [17].

The clustered distributions for some probes had been addressed previously with lesser detail [13, 14], but the Daca-Roszak study underscored the need to better characterize these probes more broadly. In that endeavor, several challenges need to be addressed. First, it would be useful to have a method to efficiently find these probes in a particular data set, rather than relying on reference data; the Daca-Roszak probe-by-probe approach [17] is not feasible for empirically assessing all 450k probes. Second, it will be useful to attribute methylation clusters to underlying genetic polymorphism where appropriate, again in a study population-specific manner. Assessing this phenomenon will require not only a careful consideration of C and G site SNPs as done previously [17], but a similarly precise examination of SBE (for Type I probes) and probe-mapping SNPs as well. Finally, it is crucial to develop a standard practice for the use or accommodation of these probes in an EWAS pipeline, since this will ultimately impact the interpretation of any DNAm association.

In our exploration of 450K data, we first noticed such clustered distributions by the “gap” pattern apparent when methylation signals per mode clustered into non-overlapping groups. In this paper, we present a method, termed ‘gap hunting’ to identify 450k probes that result in such a distributional “gap”. Identification of 450k probes with clustered methylation values using the empirical approach we propose here overcomes previous limitations with other probe removal approaches [16, 17] because it examines all measured sites, is specific to the study sample rather than relying on external annotation, which may or may not be appropriate for a particular population, and provides flexibility for the user to determine whether flagging or filtering these probes is appropriate based on their particular study design. We apply this method in a peripheral blood DNA study population from the Study to Explore Early Development, report the extent of gap signals in this dataset, and explore the sources of gap signals, with a particular focus on the various kinds (C/G sites, SBE, and probe-mapping) of SNPs and the mechanism by which they can result in a gap signal. We also describe various cases in which a probe may be affected by a SNP, but not result in a gap signal. We explore different applications of gap signals, such as their utility to be used for population stratification adjustment and their potential to enhance association analysis through discovery of methylation sites mediating genetic signal. Finally, we describe our recommendations for the role of gap hunting in the current 450k analysis pipeline.

## Results

### Identification of gap signals using gap hunting

We developed a simple, computationally fast algorithm, called ‘gap hunting’, to flag probes on the 450k exhibiting distributions of percent DNAm that cluster into discrete groups. We applied this to DNAm data from 590 whole-blood derived samples from the Study to Explore Early Development I (SEED I) at various combinations of user-supplied arguments to this procedure termed ‘threshold’ and ‘outCutoff’ (**Additional File 1**, see Methods for a detailed description of the approach and these arguments). Of the 473,864 autosomal probes we measured in SEED I participants on the 450k, we identified 11,007 (2.3%) with clustered distributions of DNAm values which we term ‘gap signals’. These results were generated using a ‘threshold’ value of 0.05 and an ‘outCutoff’ value of 0.01; the following analyses were all conducted considering these arguments and this list of gap signals. The vast majority of gap signals were composed of 2 or 3 clusters of DNAm values (**Additional Files 2 and 3**). For example, the distribution of percent DNAm for cg01802772 clusters into 3 distinct groups (**Figure 1**, top panel). Using genotyping data, available from the same SEED individuals, we found that these 3 methylation clusters correspond to genotype for SNP rs299872; this SNP is located at the interrogated C site (**Figure 1**, top panel). For this particular probe, we also queried the dbSNP138 database and found that a C/T SNP is annotated as overlapping the interrogated C site (**Figure 1**, bottom panel).

**Figure 1:**
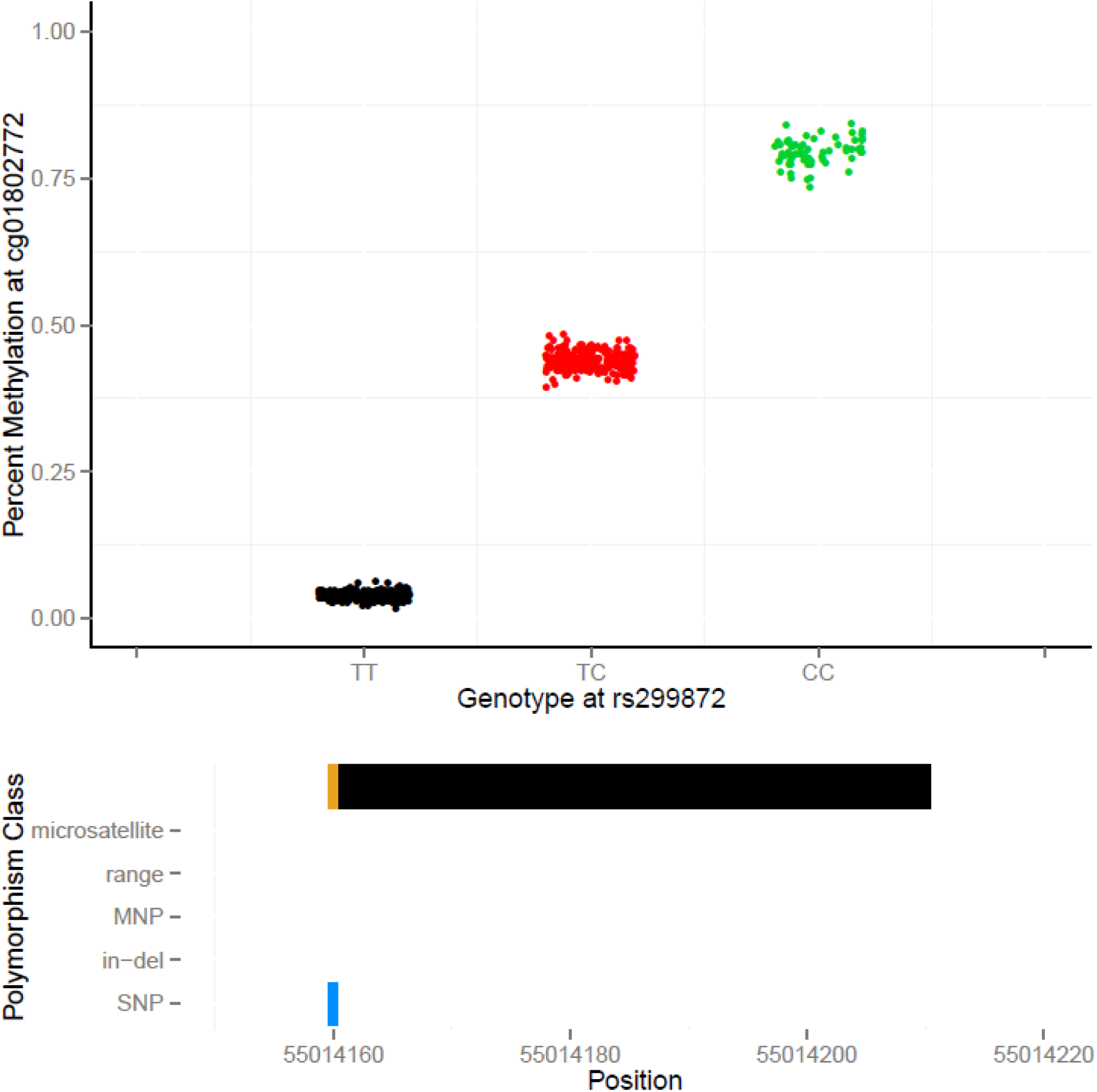
An example of a gap signal detected in SEED at cg01802772 via gap hunting. Top panel: Gap hunting identified groups are shown in black, red, and green and correspond to measured SEED genotypes TT, TC, and CC, respectively at rs299872. Bottom panel: Depiction of variant locations relative to probe orientation. Blue color denotes the single base extension site which also corresponds to the interrogated CpG site for this probe type (Type II); black color denotes 50 bp probe length. Y-axis lists variants present in the dbSNP138 database with a frequency greater than 0.5% and validated in more than 200 people.

Based on our initial gap signal observations, we decided to perform an in-depth analysis of all 11,007 gap signals to characterize the underlying source of these DNAm distributions. Using paired genotype (GWAS) and methylation data, we found that 5,453 gap probes (49.5%) contain a SNP from our SEED GWAS dataset, and thus direct evidence for SNP influence. Of the remaining gap probes, 3,746 (34.0%) have an (dbSNP13) annotated SNP, in-del, microsatellite, or multi-nucleotide polymorphism or map to a University of California, Santa Cruz (UCSC)-annotated repeat (and thus have indirect evidence for SNP/variant influence), and 1,808 gap probes (16.4%) did not contain a SNP from our SEED GWAS dataset and also were not annotated as containing a variant from the dbSNP138 database or a UCSC repeat. Given the large proportion of SNP-associated gap probes, we first sought to examine the role of SNPs in producing gap signals. Our approach to understanding the role of SNPs in producing gap signals consisted of two main elements. First, we theoretically conceptualized how various types of SNPs located at different locations in the probe, including the measured C and corresponding G loci, the SBE site, as well as elsewhere in the probe would affect 450k signal based on our knowledge of the measurement chemistry. Second, we performed empirical analyses using our joint GWAS and 450k DNAm data from SEED. We also examined the remaining ~16% of gap probes that do not have an associated SNP or variant, according to the SEED GWAS data and reference annotations.

### Predicted influence of SNPs on 450k DNAm signals

Based on the underlying 450k probe chemistry, we predicted how SNPs influence 450k signal. Our predictions are summarized in **Figure 2**. We first predicted the influence on signal of nucleotide changes for SNPs that overlap the C nucleotide of the measured locus. For Type I forward strand probes containing a T/C SNP at the interrogated C site, we predict the no signal in the methylated channel and signal in the unmethylated channel. Thus, the signal readout would be the same as for an unmethylated CpG state. For all other possible SNPs, including A/C and C/G, we would expect no signal to be reported by either the methylated or unmethylated channels for Type I forward strand probes, resulting in no overall signal; these are likely to be detected as failed probes. We predict Type II forward strand probes containing a T/C or A/C SNP at the interrogated C site to result in cyanin5 (Cy5) signal, thus, mimicking an unmethylated state. Type II forward strand probes containing a C/G SNP are predicted to result in cyanin3 (Cy3) signal, thus, mimicking a methylated state. For all reverse strand probes, including Type I and II, any SNP at the interrogated C position results in no signal from either channel, and these probes are also likely to be detected as failures.

**Figure 2:**
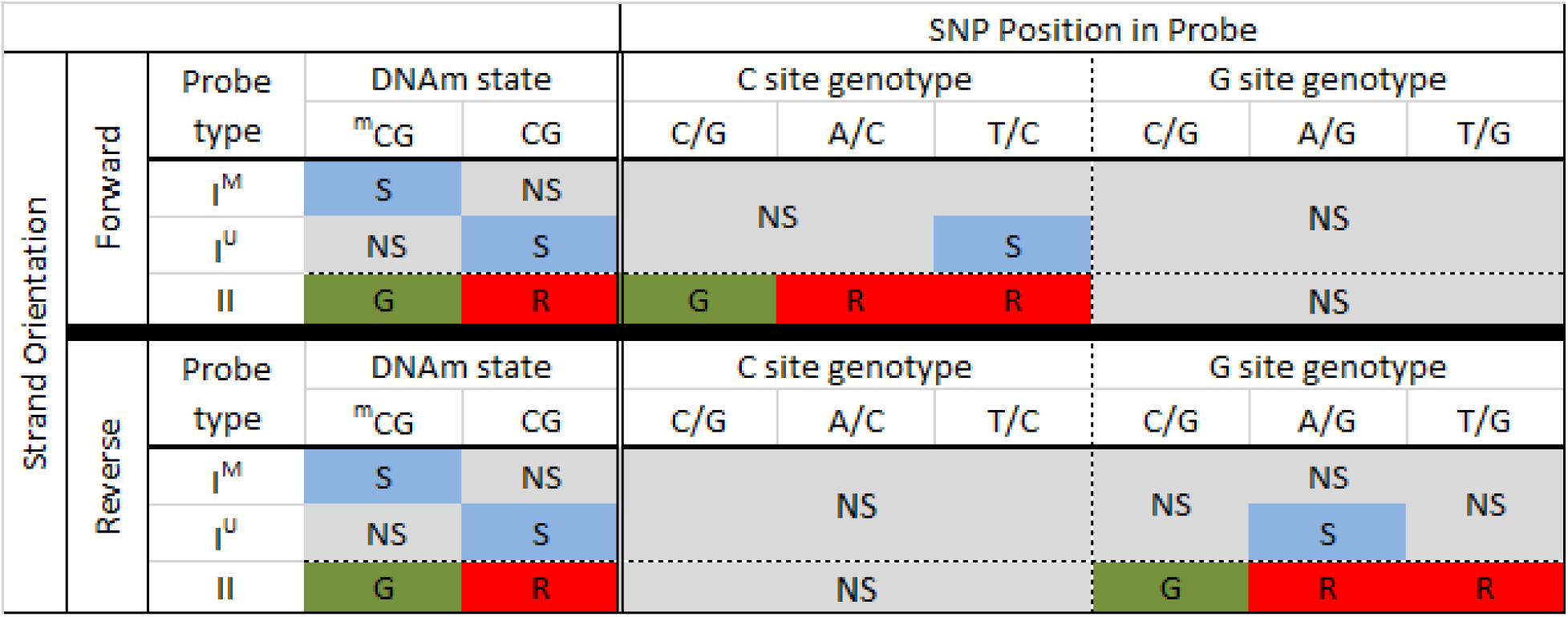
Predicted 450k signal for SNPs present at the interrogated CpG site. On the left, in the ‘DNAm state’ column, we show the expected signal for methylated and unmethylated CpG states, when no SNP is present, for both Type I and II probe designs. Middle (‘C site SNP’) and right columns (‘G site SNP’) provide expected signals for SNPs in the C and G nucleotide positions, respectively. For all columns, S denotes signal, NS denotes no signal, G and R denote red and green channel signals, respectively. ^m^CG represents methylated cytosine. I^M^ and I^U^ denote probe design type I methylated and unmethylated probe types, respectively; II denotes probe type II. For type I design, methylated probes fluoresce and unmethylated probes yield no signal when methylation is present. The type II design fluoresces in the green and red channels for methylated and unmethylated states, respectively. For forward strand interrogated CpG sites (top), a C to G SNP mimics the methylated state; C to A and C to T SNPs mimic the unmethylated state for Type II probes but result in no signal for the Type I design. One exception is for a C to T SNP because it mimics post-bisulfite converted unmethylated Cs. G site SNPs on the forward strand produce no signal for both probe designs because they inhibit single-base extension. Reverse strand probes (bottom), are defined relative to the top strand, so the expected signal scenarios are the converse of what is expected for the forward strand (i.e. G site with some signal, C site with comprehensively no signal).

Next we predicted the influence of nucleotide changes for SNPs that overlap the G nucleotide of the measured CpG site on signal detection (**Figure 2**). For Type I reverse strand probes, we predict an A/G SNP will result in a signal in the unmethylated channel and no signal in the methylated channel, resulting in a similar methylation readout as an unmethylated cytosine nucleotide. Other SNPs, including C/G and T/G, are predicted to result in no detectable signal in either the methylated or unmethylated channels. For Type II reverse strand probes, we would expect a C/G SNP present at the interrogated G site to result in a green signal, the readout for probes with these SNPs thus matches the readout for a methylated state. The presence of an A/G or T/G SNP at the G nucleotide position should result in detection of a red signal; the readout from probes with these SNPs would match the readout for an unmethylated states. Forward strand probes with any type of G-site SNP are not predicted to mask methylation states, but instead they should produce no overall signal.

Finally, we predict the influence of SNPs that overlap the SBE site on signal. For Type II probes, the SBE site overlaps with the interrogated C site; therefore, the influence of SNPs is the same as for C site SNPs, as described above and shown in **Figure 2**. In the Type I probe design, the SBE site is immediately adjacent to the interrogated C site; it is one base upstream of forward strand probes and one base downstream for reverse strand probes. The Illumina detection software is programmed to read a pre-defined color channel, which is based on the nucleotide that is expected to be incorporated (defined *a priori* using a reference sequence). For example, if the base upstream of the interrogated C site is defined as an A nucleotide in the reference sequence, the detection software will only detect signal in the red channel and will not query for a signal in the green channel. Therefore, any SBE-associated SNPs will result in a loss of signal when the incorporated nucleotide is tagged in the opposite color to that dictated by the reference sequence. As shown in **Figure 3**, C/G, A/G and T/G genotypes at an SBE associated SNP will result in loss of signal on forward strand probes. Note that the fluorescence still occurs upon SBE, but the software does not read the signal because it is in a different color channel than what is expected, based on the pre-defined reference sequence. Similarly, C/G, A/C, and T/C SNPs at SBE sites for reverse strand probes are predicted to result in loss of signal (**Figure 3**). Several SBE associated SNPs are also predicted to have no impact on the methylation readout. These include T/A, A/C, and T/C variants for forward strand probes and T/A, A/G, and T/G variants for reverse strand probes (**Figure 3**).

**Figure 3:**
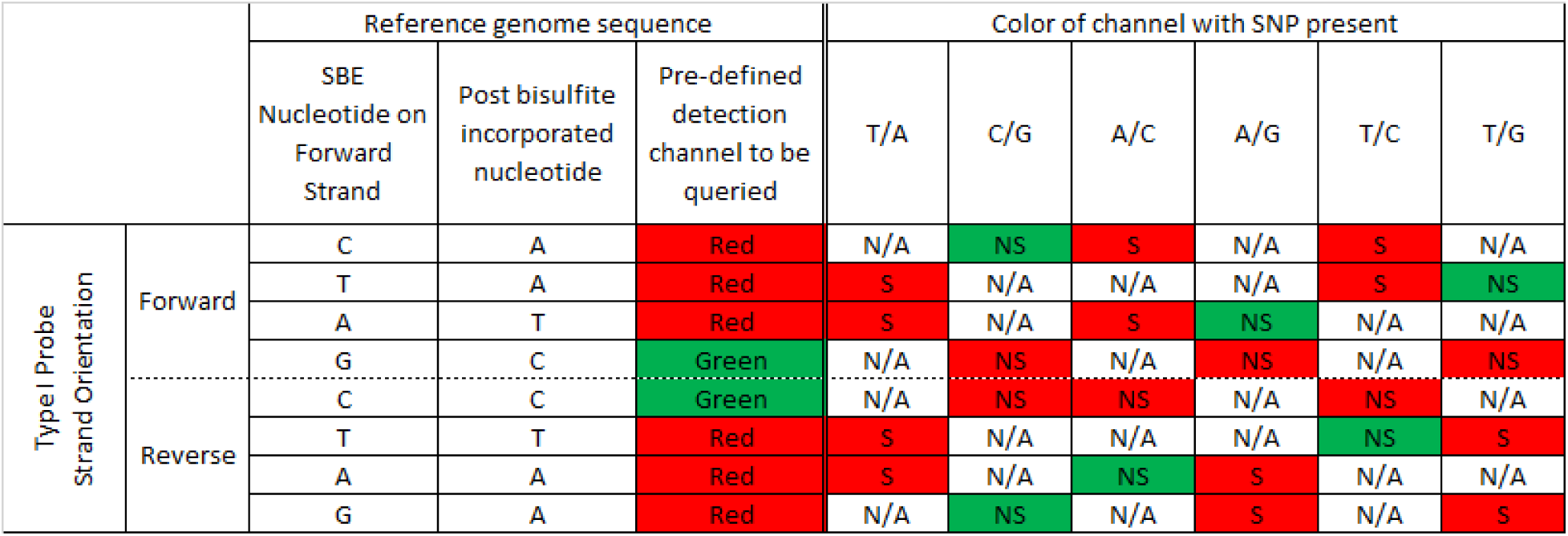
Predicted type I probe signal for individuals with a SBE site-associated SNP. For Type I probes, the SBE is located 1 bp upstream of the C site for interrogations on the forward strand, and 2 bp downstream of the C site for interrogations on the reverse strand (defining the C site location using the forward strand). Enumerating signal expectations requires consideration of bisulfite conversion, complementary bases, the expected color channel for fluorescence, and if those latter two factors change in the presence of SNP. Of note is that C and G bases are labeled to fluoresce green signal while A and T bases are labeled to fluoresce red signal (hence the existence of ‘Type I Red’ and ‘Type I Green’ probes). For example, consider a forward strand type I probe with a C nucleotide at the SBE position, based on a reference genome sequence (top row). After bisulfite conversion this base will change to a T, the complementary SBE base is an A, which fluoresces in the red channel. If instead of a C there is a G at the SBE due to a C/G SNP, the SBE incorporated nucleotide would be a C and fluoresce in the green channel. Because the software is programmed to read only the red channel, no fluorescent signal will be detected when a G SNP is present. Inferring the scenarios for interrogating a CpG site on the reverse strand requires similar reasoning but with the added consideration of complementary bases. Abbreviations: N/A – not applicable (that SNP cannot exist there), S – signal, NS – no signal.

SNPs can also occur elsewhere in the probe length, however, it is less straightforward to develop theoretical rules or principles guiding how these may affect probe signal. Similarly, it is unclear how probes with multiple SNPs may behave with respect to methylation signal. Therefore, we do not provide a theoretical framework for these types of probes, but instead provide the results from our empirical analyses below.

### Empirical evidence shows SNPs at the interrogated CpG site are related to gap signals

We performed empirical analyses to determine the relationship between DNAm levels reported by 450k and SNPs present at the CpG site using our unified SEED DNAm and genotyping data and compared them to our theoretical expectations, shown in **Figure 2.** We identified all of the 450k probes with a measured or imputed SNP present at the interrogated C or corresponding G loci in our SEED sample (n=5,129) (**Additional File 4)**. To ensure we were only assessing the influence of our SEED SNPs at CpG sites, we limited our analysis to include only probes with a SNP at the CpG site itself and not elsewhere in the probe length. We found that our empirical SNP results coincided with our predicted results for the SNP scenarios shown in **Figure 2 (Additional File 5)**. For example, we observed a positive correlation between percent DNAm and dosage of the G allele across the set of 94 probes, including 23 Type I and 71 Type II probes, containing a C/G SNP at the interrogated C locus (**Figure 4A** and **Additional File 4**). This appears to be a direct consequence of the positive correlation with methylated probe signal and negative correlation with unmethylated probe signal (**Figure 4B-C**). This observation coincides with our prediction for this scenario (**Figure 2**) because the addition of the non-reference (G) nucleotide is expected to increase methylated (green) signal at the expense of unmethylated (red) signal. To better conceptualize the effect of a SNP on the total produced signal, i.e. combined methylated and unmethylated signals, we computed a copy number metric (see Methods) and found, in general, it decreased with dosage of the G allele (**Figure 4D**). However, the mean copy number metric of the heterozygous group does not lie exactly intermediate between the two homozygous groups, thus highlighting the importance of also considering the methylation state in the interpretation of SNP-influenced 450k probes. For example, in **Figure 1** (top panel), individuals with the ‘TT’ genotype have low methylation values because of their low ratios of methylated to unmethylated intensities dictated by their T allele. Individuals containing one or two copies of the C allele at this SNP can have varying degrees of methylation. In the example shown in **Figure 1**, the C alleles are completely methylated for all samples, resulting in discrete DNAm groups. If however, the C alleles were unmethylated, the groups would be largely indistinguishable and form one cluster instead of three. The lack of an explicitly intermediate mean in the heterozygous group for the copy number metric, then, is a consequence of the heterogeneity in methylation at the ‘C’ base at these sites and heterogeneity amongst samples in their methylation state. **Additional File 5** contains plots for the remaining SNP scenarios delimited in **Figure 2**, and all showed similar relationships.

**Figure 4:**
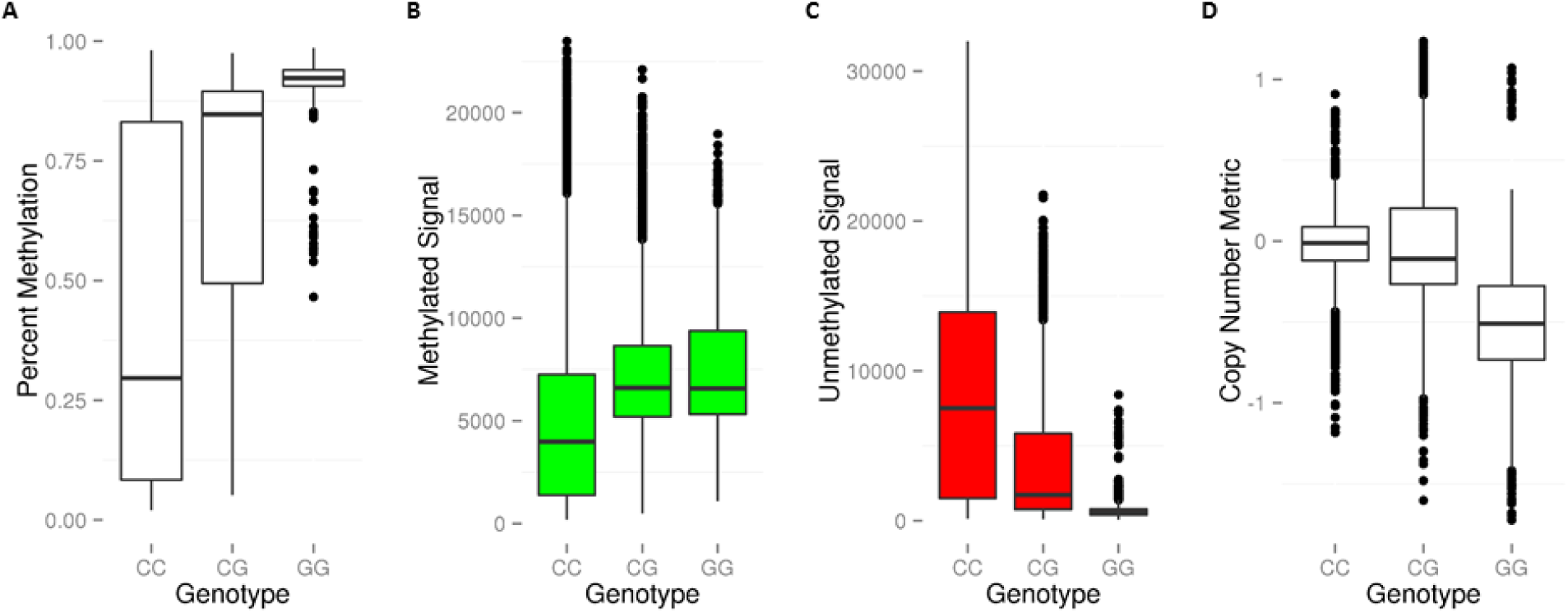
The influence of a C/G SNP located at the interrogated cytosine on reported methylation signal in Type II forward strand probes. (**A**) Percent methylation vs. genotype plot shows a positive correlation between percent methylation and dosage of the G allele. (**B**) Methylated signal vs genotype plot shows a positive correlation between methylated signal and dosage of the G allele. (**C**) Unmethylated signal vs genotype plot shows a negative correlation between methylated signal and dosage of the G allele (**D**) Copy number metric vs genotype plot shows a negative correlation between copy number and dosage of the G allele.

### Empirical evidence shows SBE site SNPs are related to gap signals

We identified all of the 450k probes in SEED with a measured or imputed SNP located at the SBE site (n=118) (**Additional File 4**). We specifically limited our analyses to probes that contained an SBE-associated SNP exclusively, i.e. there were no SNPs elsewhere in the probe. We found that, overall, our empirical results correspond to our predicted signal for the SNP scenarios shown in **Figure 3 (Additional File 6)**. For example, we observed an inverse relationship between dosage of the T allele and overall signal across all probes (n=2) that have a T/G SNP at the SBE site, where G is the *a priori* defined base at the SBE according to the genome reference sequence (**Figure 5**). This observation coincides with our prediction for this scenario (**Figure 3**) because a SNP changing the nucleotide at the SBE position from ‘G’ (detected in the green channel) to ‘T’ (detected in the red channel) should result in no signal because the software is programmed, *a priori* based on reference genome sequence, to report methylation solely as a function of the signal being generated in the green channel. Note that similar to CpG associated SNPs, the mean copy number metric of the heterozygous group does not lie exactly intermediate between the two homozygous groups. This is likely reflecting the heterogeneity in DNA methylation across CpG sites and samples. Overall, our findings using SEED measured and imputed genotypes are consistent with our predictions shown in **Figure 3.** However, in certain cases the relationship is less clear. There are a number of potential explanations for non-linear relationships. First, since the overall signal is a measure of both the ability to detect signal, which as we’ve shown above can be influenced by SBE site SNPs, and the actual methylation state itself, it is possible that deviations from the expected relationship are related to actual differences in DNAm. These DNAm influences may be exacerbated by the relatively small number of probes examined in each scenario shown in **Figure 3** (all scenarios have ≤ 17 probes and some scenarios have ≤ 7 probes; see **Additional File 4**). It is also possible that these non-linear genotype signal shifts could be related to uncertainty around imputed genotypes.

**Figure 5:**
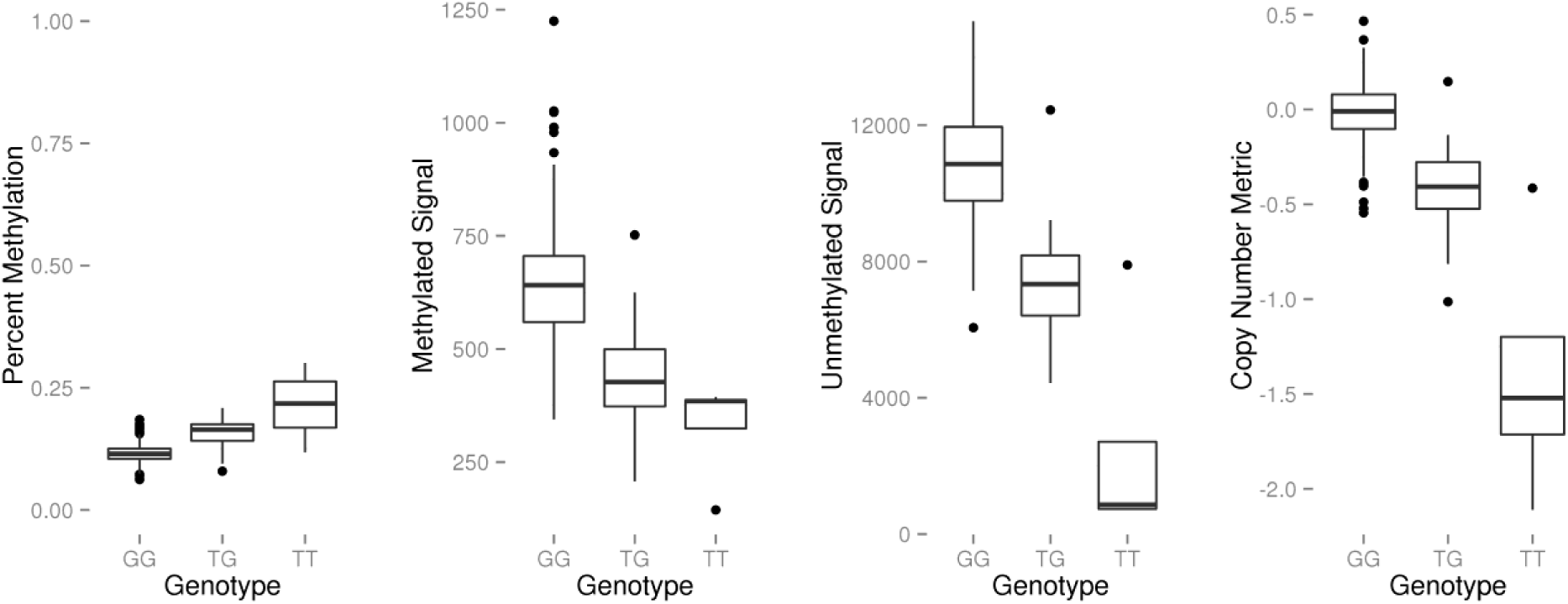
The effect of a G/T SNP at the SBE site of Type I probes on percent methylation, methylated signal, unmethylated signal, and a copy number metric. Percent methylation (beta value), methylated signal, unmethylated signal, and a copy number metric plotted against genotype for Type I probes interrogating a CpG site on the forward strand, when the G is the reference genotype. Information was collected across 2 probes. There is an inverse association between dosage of the T allele and signal produced, as predicted in Figure 3.

### Probe SNPs up to 20 base pairs from the CpG site are associated with gap signals

Next we sought to specifically assess the relationship between the gap signals we detected via gap hunting and probe SNPs using our unified SEED GWAS and DNAm data. We identified all of the 450k probes in SEED with a measured or imputed SNP located in the probe, excluding those with SNPs at the SBE or CpG sites (n=33,317). We limited our analysis to probes that contained a single SNP to determine the relationship between SNP distance to the interrogated C site and gap signal. The probes were binned by SNP distance to the interrogated C site and samples were grouped by genotype: homozygous for the reference allele, heterozygous, and minor allele homozygous. When we plotted the signal intensities, for both the methylated and unmethylated channels, which represent the mean intensity for all probes with a SNP at that particular distance to C-site, we observed differences in mean signal intensities, and inter-quartile ranges (25^th^ – 75^th^ percentiles), between heterozygotes and homozygotes that were consistent with allelic dosage (**Figure 6**). For the Type II probe design, mean intensity differences between the genotype groups are observed up to a SNP distance of about 7-8 base pairs (bp) from the interrogated C-site. We also observed that these probe SNP-related differences in signal intensity are less pronounced in the methylated channel compared to the unmethylated channel, where differences in intensity can persist for up to an approximately 20 bp distance. Thus, the unmethylated signal channel appears to be less robust to probe SNPs. Type I probes exhibit a similar behavior, but appear to show greater differences in signal intensity with SNPs and across larger probe distances (**Additional File 7**). One explanation for this behavior could be that Type I probes are more susceptible to probe SNPs because they were designed under the assumption that the interrogated CpG site and any CpG sites throughout the remainder of the probe length have the same methylation state (**Additional File 7**).

**Figure 6:**
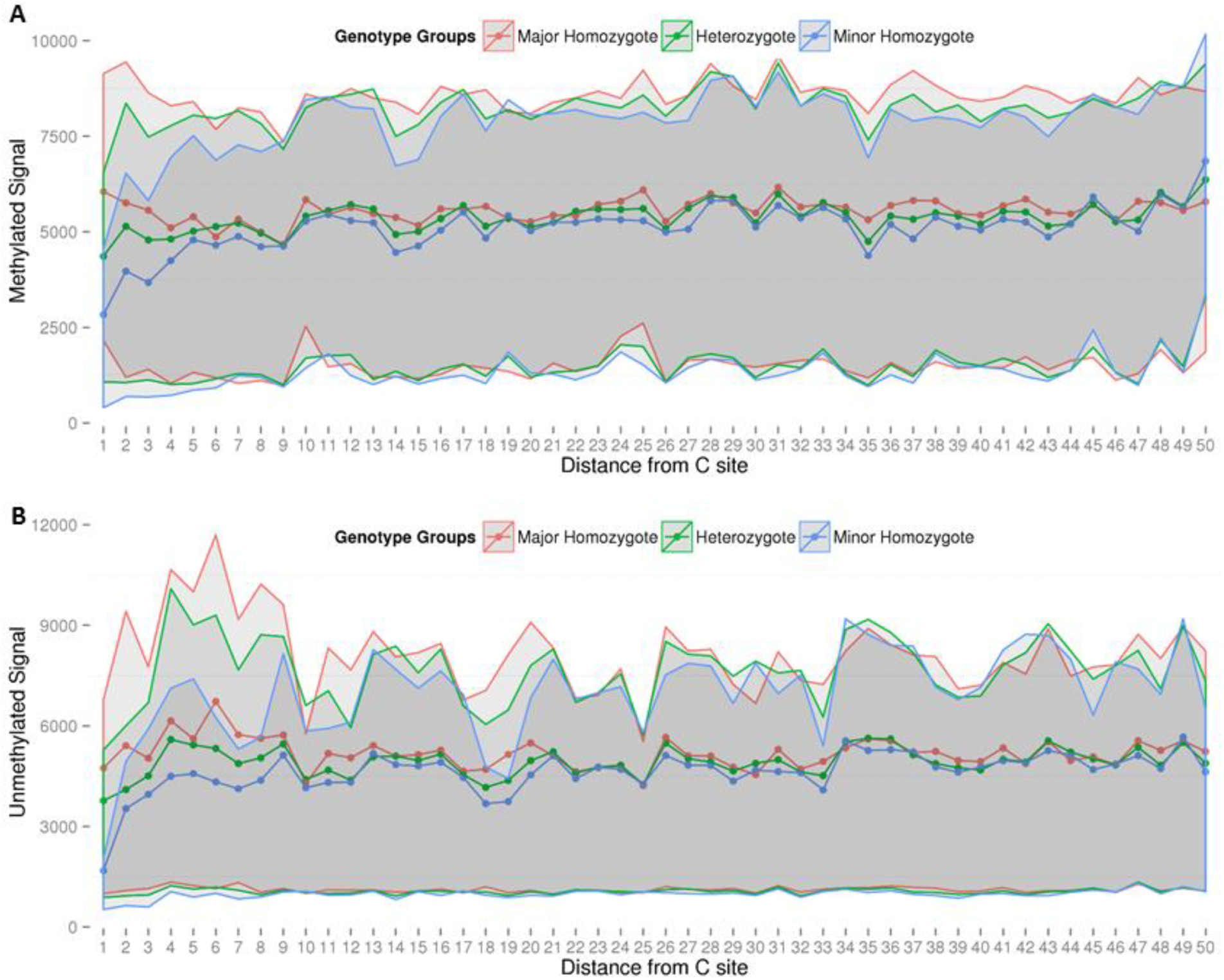
The effect of probe SNPs on methylated signal and unmethylated signal in Type II probes. We isolated specific probes that met the following conditions: it contained a measured SNP in the 50bp probe length outside of the C, G and/or SBE sites, and it contained only a single SNP in the probe length. The probes that met our criteria varied in distance from 1-50 base pairs from the interrogated CpG site. At each distance value, we plotted the mean (shown by dotted lines) and inter-quartile range (greyed area) of the people who were homozygous for the reference allele (shown in red), heterozygous (shown in green) or homozygous for the minor allele (shown in blue). Lack of signal concordance across these 3 groups indicates stronger SNP influences on signal. For both methylated (**Panel A**) and unmethylated signals (**Panel B**), polymorphisms closer to the C site show stronger influences on signal. The influence is strongest up to approximately 10 bp but is observed up to roughly 20 bp from the measured C-site.

### SNP-affected probes do not always result in gap signals

Our analyses above focus on identifying potential sources of gap signals and show that SNPs can lead to gap signals. Therefore, we also wanted to determine whether probe-associated SNPs always lead to gap signals. We found that not all polymorphism-affected probes result in gap signals (**Additional File 4**). There are 3 main classes of beta distributions in which a probe may be affected by a SNP, but not result in a ‘gap-like’ distribution (**Figure 7**). The first occurs when there is a correlation between percent methylation and genotype, but no discrete clusters are observed (**Figure 7A**). The second occurs when there are outlier signals, i.e. samples. The gap hunting algorithm was designed to exclude probes from the gap signal list if they likely contained an outlier sample. As a result if the smallest group of samples driving SNP-related gaps is less than the proportion of samples determined by the ‘outCutoff’ argument, these probes will not be flagged as gap signal probes. **Figure 7B** illustrates this point; it shows that at cg15013523, gap hunting would not identify the group with the ‘TT’ genotype as a discrete cluster, i.e. gap signal, because it is comprised of a single sample. These types of probes could be identified as a gap signal if the option to retain ‘outlier-driven’ probes is selected. Finally, beta distributions with an associated SNP in the probe may show no DNAm variability at the site or no correlation with genotype and, therefore, will not result in a ‘gap-like’ distribution (**Figure 7C**); this lack of clear genotype correlation was also observed by Daca-Roszak *et al* [17] and referred to as a ‘cloud-like’ distribution. Therefore the potential for a polymorphism-affected probe to be classified as a gap signal is related both to the presence of discrete separation in groups, as well as the overall methylation state at the site.

**Figure 7:**
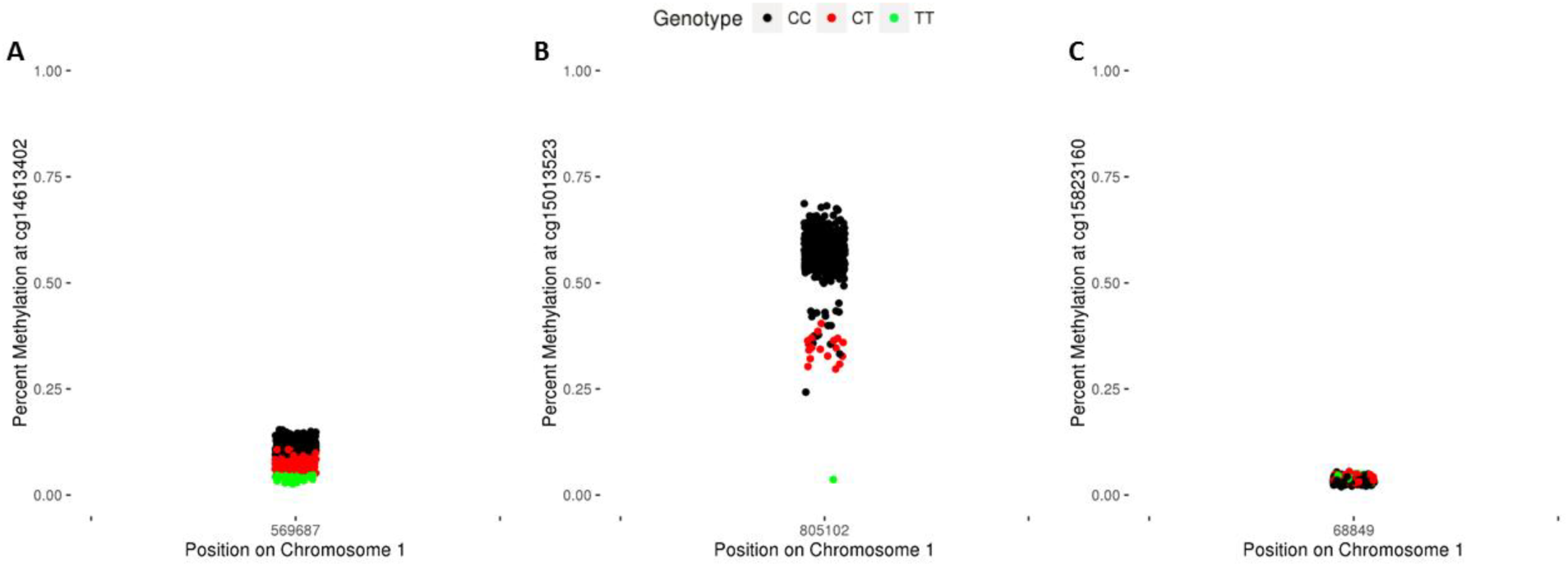
Examples of probes with a polymorphism that do not result in a gap signal. Most probes that overlap with SEED SNPs are not classified as gap signals. These probes can generally be grouped into 3 categories: **Panel A**: In SEED, cg14613402 overlaps with a C/T SNP at the interrogated C site and displays a negative correlation with dosage of the T allele. However, a discrete difference in the groups is not achieved. **Panel B**: cg15012523 overlaps with a C/T SNP at the interrogated C site and also displays a negative correlation with dosage of the T allele. Here, a discrete difference does existence between the TT genotype and others and thus would be identified via gap hunting; it would be classified as an outlier-driven signal with the default algorithm arguments, however (see Methods). **Panel C**: cg15283160 overlaps with a C/T SNP at the interrogated C site but displays no variability in beta value.

### Approximately 16% of gap signals identified in SEED cannot be attributed to an underlying SNP

Finally, among all autosomal probes, we compared the standard deviation distribution between gap and non-gap probes, both with and without an associated SNPs, to better characterize gap signals that could not be attributed to an underlying SNP. The 6 mutually exclusive classes of probes we examined include (1) non-gaps with measured or imputed SEED SNPs, (2) non-gaps with annotated variants or repeat elements, (3) non-gaps with no associated variants, (4) gaps with measured or imputed SEED SNPs, (5) gaps with annotated variants or repeat elements, and (6) gaps with no associated variants. As shown in **Figure 8**, all non-gap probe distributions, including those with and without an associated SNP, are highly overlapping for both the Type I and Type II designs, suggesting that the majority of non-gap probes have no or low variability in DNAm values similar to the example in **Figure 7C**. The gap probe distributions are distinct from the non-gap distributions and show interesting within-group differences (**Figure 8**). The gap signals with reference database annotated SNPs exhibit a higher proportion of probes with larger standard deviations than those with SEED measured or imputed SNPs. This is likely due to higher minor allele frequencies of annotated SNPs, generally, compared to the minor allele frequencies of the SEED SNPs (**Additional File 8**). Another interesting feature of the overall standard deviations is the distinct curve of the 1,808 gap signals that lacked any of a measured/imputed SNP, reference database annotated variant, or a UCSC repeat in the probe. As clearly seen in the Type II probe design, there is a high proportion of gap probes without an associated SNP or annotated variant/repeat at low standard deviation values, relative to gap probes containing SNPs (**Figure 8**). We also show that gap probes without a SNP or annotated variant/repeat tend to have a higher proportion of 2-cluster probes than gap probes with a SNP (**Additional File 9**).

**Figure 8:**
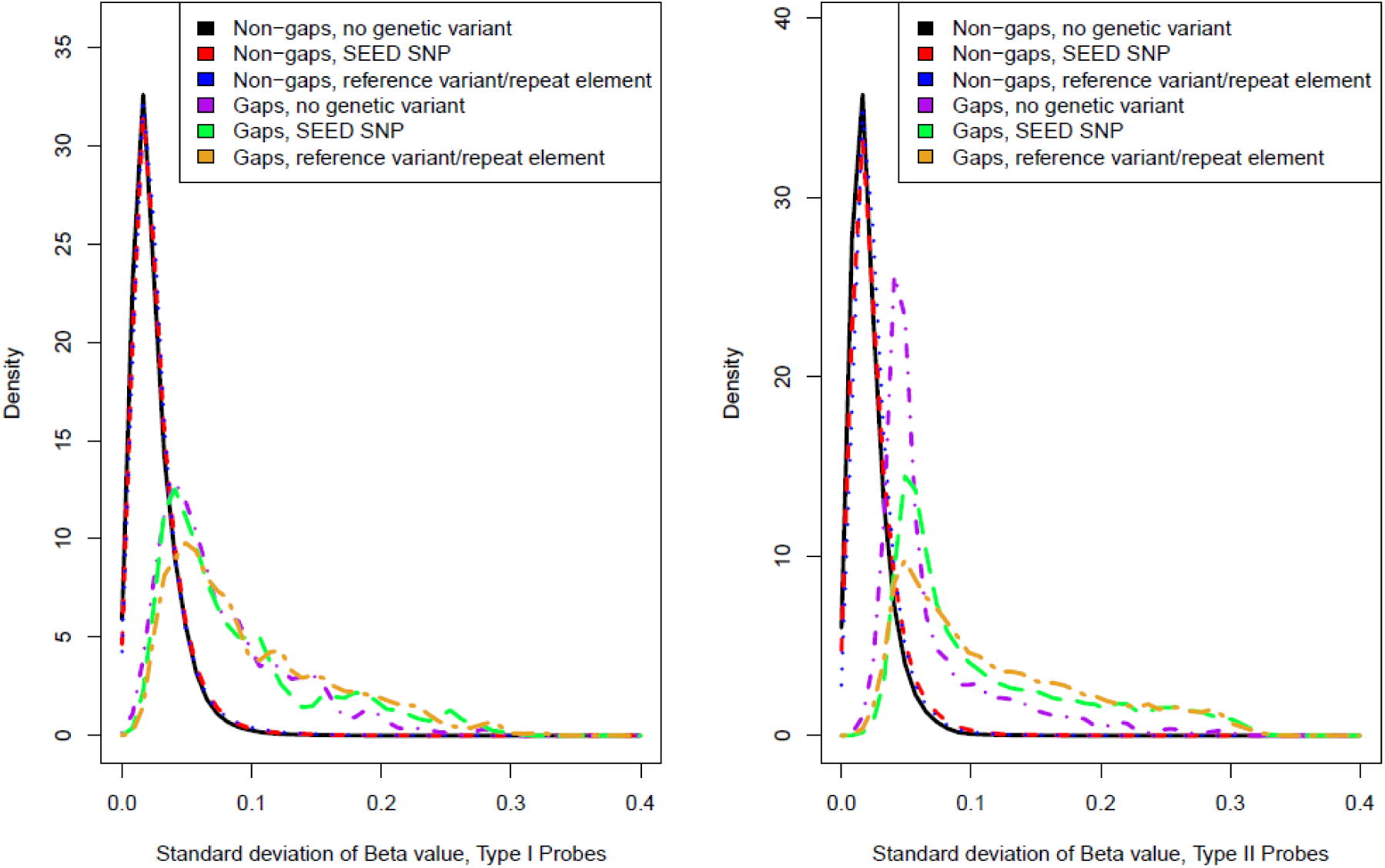
Distributions of standard deviations among 6 categories of 450K probes. All autosomal probes (n = 473,864) were classified into one of six groups: (1) non-gap probes that lack a SEED SNP, dbSNP-annotated polymorphism, or UCSC-annotated repeat that map to the probe (n = 301,590; shown in black), (2) non-gap probes with at least one SEED SNP present in the probe (n = 62,005; shown in red), (3) non-gap probes that do not contain a SEED SNP but do have an annotated variant as indicated by the dbSNP138 database or map to a UCSC-annotated repeat (n = 99,262; shown in blue), (4) gap probes that lack a SEED SNP, dbSNP-annotated polymorphism, or UCSC-annotated repeat that map to the probe (n = 1,808; shown in purple), (5) gap probes with at least one SEED SNP present in the probe (n = 5,453; shown in green), (6) gap probes that do not contain a SEED SNP but do have an annotated SNP as indicated by the dbSNP138 database or map to a UCSC-annotated repeat (n = 3,746; shown in orange). The 3 non-gap probe distributions are distinct from the gap probe distribution but show some overlap; suggesting some probes with ‘gap-like’ distributions are not captured by gap hunting (also see **Figure 7** for explanation). The gap probe distribution for those probes with annotated SNPs (green and orange) has a slightly higher area under the curve at higher standard deviation values (especially for the Type II design), which is likely due to the generally higher allele frequencies for the annotated SNPs compared to the measured SNPs (see **Additional File 8**). Gap probes lacking any probe SNPs form a distinct distribution, especially for the Type II design (purple).

We were interested in quantifying the degree to which other factors, aside from a measured/annotated SNP or annotated repeat element, could lead to gap-like behavior. For example, it is possible that some of these 2-cluster gap signals ambiguously mapped to sex chromosomes and clustered according to sex; however, we only observed sex-specific clusters in 6 (0.3%) of these probes. 161 of these 1,808 probes in total were previously defined as ambiguously mapping [12], 14 were determined to have failed via detection p-value, and 210 were found to discriminate blood cell types [18]. These 3 factors account for 379 (21.0%) unique probes of the 1,808 gap signals not attributed to an underlying genetic variant.

### Other methods to identify clustered DNAm distributions are not as robust as gap hunting

We assessed the potential for other methods to identify clustered DNAm distributions with respect to detection sensitivity and the specific types of sample clusters they identify. We tested these methods against a set of 5,000 probes made up of gap signals (which functioned as positive controls) and 5,000 probes which were not gap signals, had no measured, imputed, or annotated SNP, and had very low variability (and thus functioned as negative controls). First, using the beta values for these probes, we applied a Gaussian mixture model clustering algorithm, which selects the optimal number of clusters based on the Bayesian information criterion (BIC), and found that it had 100% sensitivity, but only 50% specificity, to distinguish between the gap and non-gap probes. Additionally, in cases where the mixture model predicted a gap signal to (correctly) have more than 1 cluster, it was only able to identify the correct number of clusters 43% of the time. We also examined the utility of the dip test, in which the null hypothesis is that the data are unimodal [19], and found the area under the receiver operating characteristic curve to be 0.73. These methods performed similarly using M-values, with a 100% sensitivity, 67% specificity and 41% correct determination of cluster number using the mixture model, and an area under the curve determined by dip test p-values of 0.73. We were then interested in examining the performance of these methods at specific scenarios (**Figure 7**) to which gap hunting was insensitive (**Additional File 10**). The mixture model approach was unable to correctly identify that there were 3 relevant clusters in any of these 3 probes, while using the dip test we could only reject the null hypothesis of uni-modality at cg14613402 (**Figure 7A**); however, this would not be the case if we used a more stringent p-value to account for multiple comparisons. Finally, these 2 alternative methods did identify probe cg01802272 (**Figure 1**) as a multimodal (dip test p-value ≈ 0) as well as having 3 discrete clusters, consistent with our gap hunting approach.

### Gap hunting can be useful in addressing population stratification in epigenome-wide association studies

After gaining an understanding of gap signal properties, we were interested in highlighting the potential utility of gap hunting in EWA studies. A recent paper by Barfield et al. demonstrated the ability of principal components (PCs) derived from probes annotated with 1000 Genomes-identified SNPs to correct for population stratification [20]. This method functions under the principal that methylation at these sites will be enriched for genotype-influenced signal, and thus serve as a suitable alternative to or surrogate for gold standard correction via genotype-derived PCs [21] in studies where genotype data is unavailable. Given the strong SNP influence on gap signals, we hypothesized that PCs derived from gap signals could be utilized in a similar manner to the Barfield method. Similar to Barfield et al, our gap signal based PCs we able to clearly separate ancestry groups (**Figure 9**). This result is expected since most (~85%) gap signals can be attributed to an empirical or reference database annotated SNP/variant, most of which are present in the 1000 Genomes Project that was used by Barfield et al. Additionally, most of the 96 probes identified by Daca-Roszak et al. because they differentiated two ancestral groups [17] exhibit ‘gap-like’ distributions.

**Figure 9:**
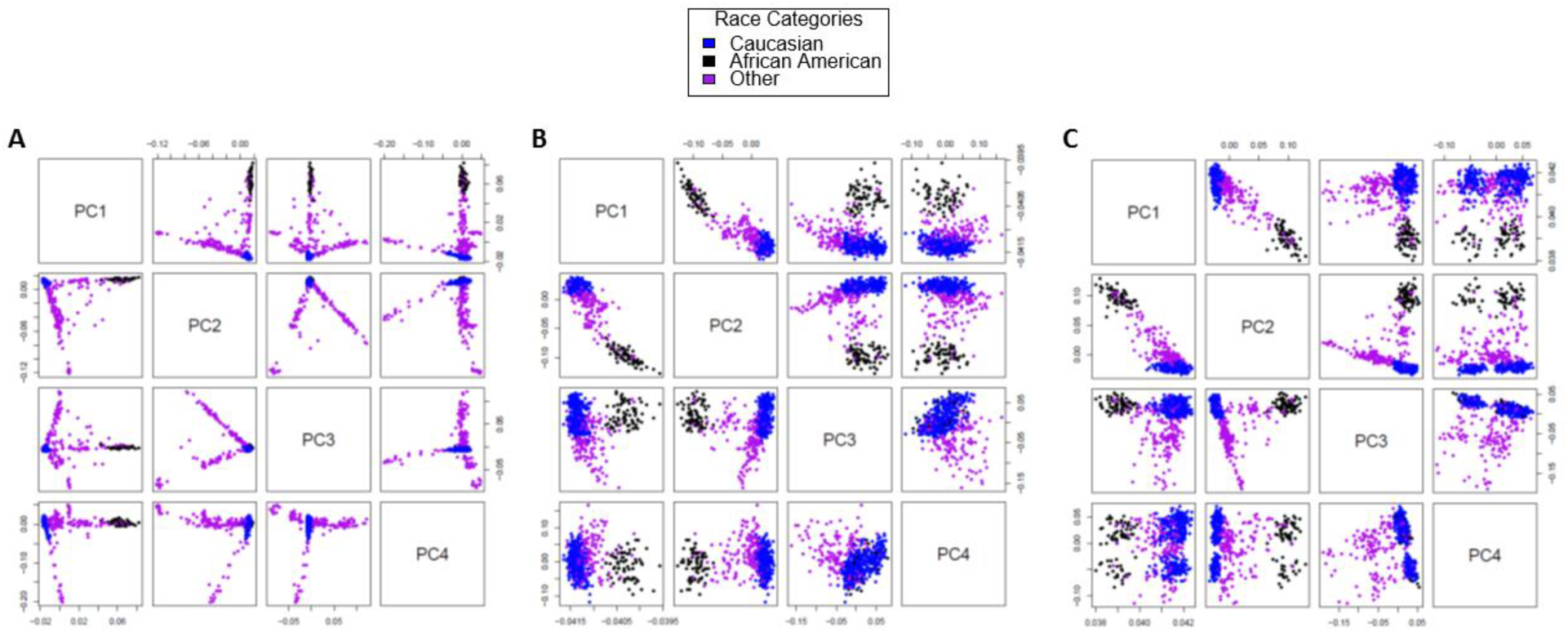
Comparison of several different methods, including gap probes, for population stratification adjustment. Points are colored according to self-reported race with Caucasian shown in blue, African American shown in black, and Other shown in purple. Each panel contains a series of plots in which the values plotted are dictated by the row (y-axis) and column (x-axis). For example the top row will plot PC 1 (y-axis) vs PCs 2, 3, and 4 (x-axis). **Panel A**: Eigenvectors generated from GWAS data using the EIGENSTRAT software [21]. **Panel B**: PCs generated from probes overlapping with 1000 Genomes-annotated SNPs (0 bp from C site option) as demonstrated by Barfield et al. [20]. **Panel C**: PCs generated from gap signals, which perform similarly to the existing methylation-based method to account for ancestry in EWA-studies show in Panel B.

### Gap signals are enriched in common EWA probe filtering strategies

One approach used in EWA studies to address multiple testing burden is to subset the dataset to only those probes that are variably methylated. We sought to define the proportion of probes in the post-variably methylated filtering dataset that had gap signals identified using gap hunting. As expected, due to our gap signals having inherently high variability, we observed gap signal probe enrichment in the filtered dataset as we increased our standard deviation threshold for filtering (**Additional File 11**). Enrichment was consistent at various percentile cutoffs of standard deviation across samples. This result emphasizes that researchers should be aware that applying filtering criteria related to probe variability can increase the proportion of gap signals.

### Common EWA probe filtering strategies that remove all SNP-associated probes may miss disease-relevant loci

Currently, most EWA studies explicitly remove polymorphism-affected probes that are *a priori* defined using a reference SNP database or in the Illumina manifest, prior to association analyses. However, based on our findings there are two main concerns with this removal approach. First, it is possible that the SNPs present in reference databases, gathered from many ancestral populations and often includes rare SNPs, may not reflect the genetic architecture among the samples examined in a particular EWA. Second, we have shown that gap signals can be influenced by SNPs and, therefore, gap signals may represent the local genetic structure underlying the interrogated CpG site; thus, they could still be biologically relevant to the outcome of interest, but should be interpreted with caution. This local genetic structure extends beyond that of the interrogated CpG site and 50-mer probe and includes the entire haplotype on which the CpG site exists. For example, cg12162195 exhibits a three-group gap signal, with three SNPs annotated in the probe body (**Figure 10**). The samples in each group represent distinct groups of haplotypes; therefore, these methylation groups serve as a surrogate for their respective collections of haplotypes. This is true for all 450k probes that are not located within recombination hotspots. Given our findings that many gap signals have a strong genetic basis underlying the observed differences in methylation, we would expect methylation values at gap probes will capture a larger degree of haplotype diversity than non-gap probes. Therefore, we propose that instead of removing reference database SNP or gap hunting-defined gap signal probes before association analyses, they be included, but flagged, and carefully investigated and interpreted after analyses should they be associated with the outcome of interest.

**Figure 10:**
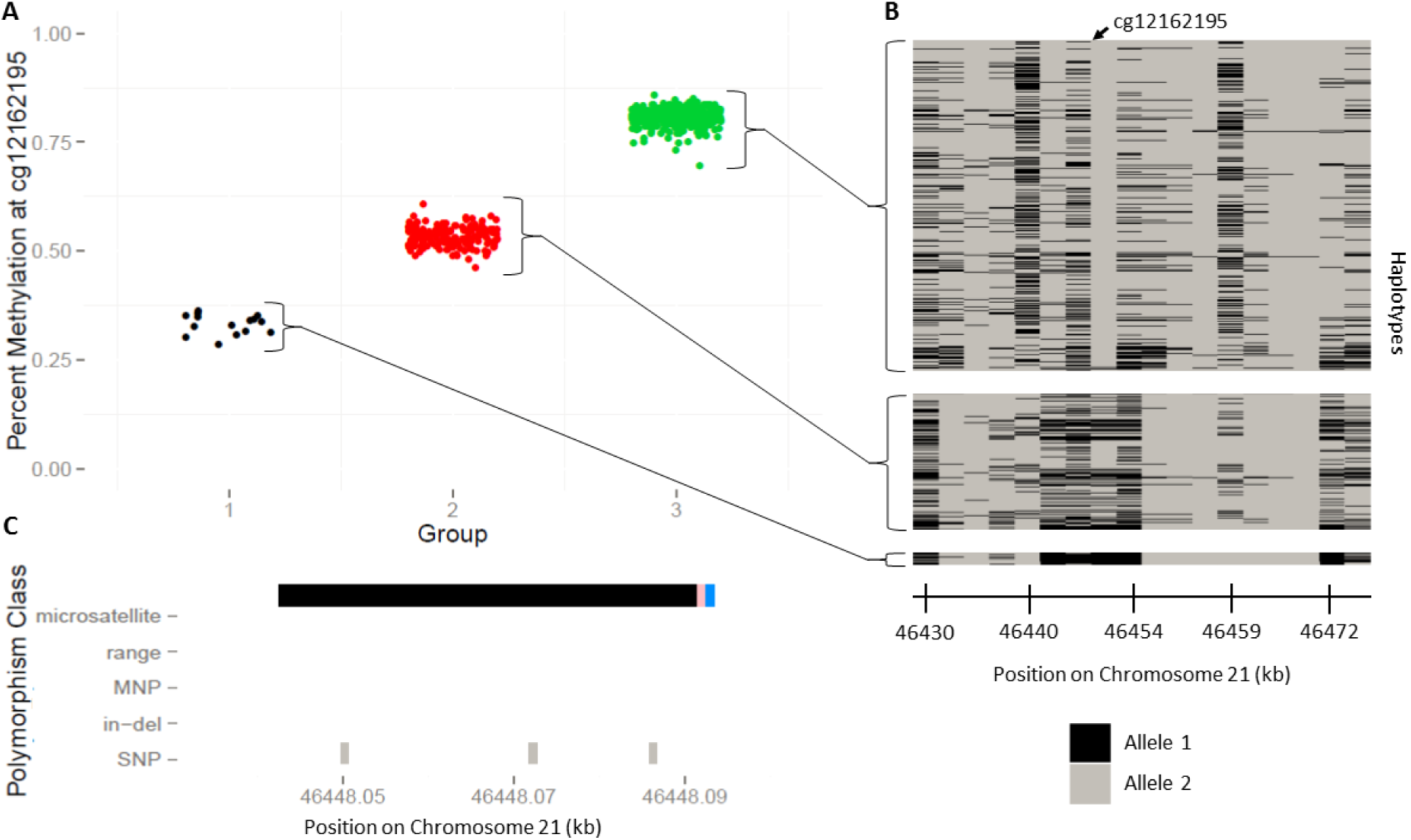
Relationship between DNA methylation (DNAm) clusters, identified by gap hunting at cg12162195, and local haplotypes among the same individuals. (A) Percent methylation at cg12162195 vs. gap hunting-defined DNAm group. (B) Individual haplotypes sorted by gap hunting-defined DNAm group. Each column represents a genotyped SNP at a specific locus across all individuals with corresponding DNAm data. Each row denotes an individual’s local haplotype for the region that contains cg12162195. There are two rows per individual, one per haplotype. The arrow at the top of the plot depicts the location of cg12162195 within the haplotype region. Gap hunting-identified groups correspond to different sets of haplotypes; these methylation groups can be used as surrogates of these haplotype groups. (C) Depiction of variant locations relative to probe orientation. Blue color indicates the single base extension site; black color denotes 450K probe; pink denotes the interrogated CpG site. Y-axis lists variants present in the dbSNP138 database with a frequency greater than 0.5% and validated in more than 200 people.

To examine the difference between our “flag” based gap hunting approach versus a “remove” reference SNP annotation based approach in downstream interpretation of EWAS, we ran our EWAS pipeline on publicly available data to evaluate the relationship between placenta methylation and newborn neurobehavior. We identified a total of 11,286 gap signals among 443,825 probes. Using our EWAS pipeline, 56 probes showed suggestive statistical significance (p < 1E-4) for infant arousal. Of these significant probes, 5 were gap signals (**Figure 11**), 15 were annotated as SNP-affected, and 3 of these were both SNP-annotated and gap identified. Thus, an analysis with gap hunting results in 56 hits, 5 of which are flagged as gaps for further investigation and interpretation. Using SNP annotation filtering without gap hunting resulted in 41 hits, 2 of which have suspicious gap-like distributions. Inclusion of all probes in EWAS, with flags for gaps via gap hunting and for SNP annotation, rather than explicitly filtering probes, allows broader consideration of biologically-relevant associations and user-specific choices regarding interpretation, rather than omitting potentially relevant findings.

**Figure 11:**
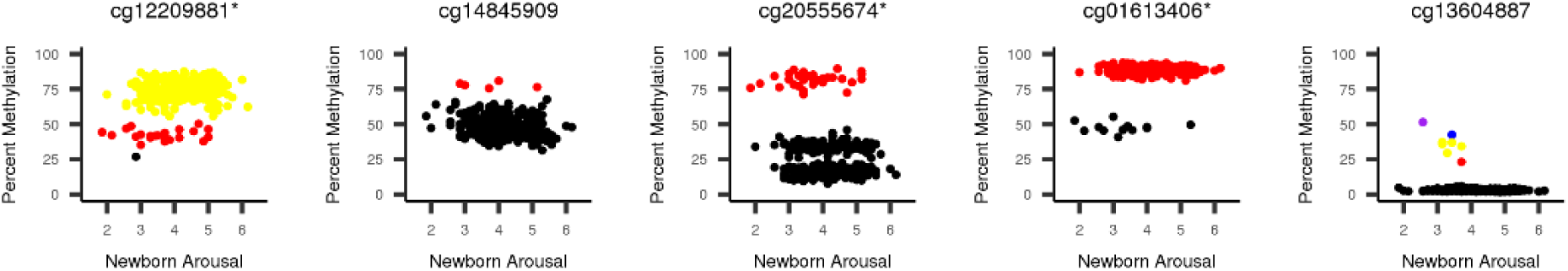
Five gap signals identified in the list of 56 probes that attained suggestive significance (p < 1E-4) with newborn arousal in a publically available dataset. There is 1 plot for each probe, with percent methylation plotted on the y-axis and newborn arousal score plotted on the x-axis. Each sample is colored by its gap hunting-identified group. The * indicates a probe that would have been filtered out via the dbSNP137 reference annotation in the *minfi* package.

## Discussion

We demonstrate a procedure called ‘gap hunting’ to identify CpGs with clustered methylation distributions and discover 11,007 ‘gap signals’ in a 450k dataset from the Study to Explore Early Development. The vast majority (~85%) can likely be attributed to an underlying SNP(s) or other variant in the C site, G site, SBE site, or elsewhere in the probe length. We document the specific mechanisms by which SNPs at the C, G, and SBE sites lead to gap signals, which involve a consideration of the type of nucleotide change occurring, as well as the probe type, DNA strand of interrogation, and overall methylation state, and demonstrated that expected effects are met using paired genotype and 450k data on the same samples. We additionally demonstrate that distance between a probe SNP and the C site of the probe is a relevant factor influencing methylation distributions. Finally, we delimit the situations in which a SNP-affected probe does not produce a gap signal, highlight a subset of gap signals that cannot be attributed to an underlying SNP, and discuss various utilities of gap signal identification in an EWAS framework. We recommend using gap hunting to ‘flag’ probes for special consideration when interpreting EWAS findings, rather than the common practice of removing probes annotated with SNPs (using reference annotations) prior to EWAS.

The focus of this manuscript was to characterize sources of clustered distributions, and highlight implications of their removal or inclusion in an EWAS framework as opposed to developing an optimized statistical method to *identify* these probes *per se*. In light of this conceptual framework, we selected a ‘threshold’ argument of 0.05 and a ‘outCutoff’ argument of 0.01 for our analyses because it balanced detecting too many probes vs. too few probes for downstream characterization and provided good face validity to capture the distributions of interest when we visually observed the data.

We recognize that changes to these parameters may influence the total number of gaps detected, indeed for our SEED dataset, the number of gap signals varied with different combinations of these parameters (**Additional File 1**), with the ‘outCutoff’ argument having the most pronounced effects at lower ‘threshold’ values. While the parameters utilized here may be a good starting point, we encourage users of the gap hunting approach to modulate this argument while understanding that reported gaps are simultaneously a function of factors such as sample size (more samples can lead to ‘denser’ DNAm distributions), disease status, or other factors that contribute to DNAm variability.

Our results highlight the importance of taking a “flag” versus removal approach. Typically, there are two main concerns motivating a removal approach: that SNP-affected probes can lead to technical failure of array measurement, and that they are redundant with genotype signal. Our work does show that C, G, and SBE site SNPs impact methylation signal, based on various factors, including bisulfite conversion and probe chemistry, as expected. This could be reinterpreted not as probe ‘failure’ of measurement, but as methylation signal that is a reliable surrogate for true underlying biology. This interpretation is, in fact, the second argument for removal often cited, but flagging for subsequent interpretation rather than removal can shed light on genome-epigenome interactions that may be relevant to the EWAS question, and would be missed via a removal strategy. These biologically illuminating relationships can relate a single SNP to a single CpG site (**Figure 1**), or can extend to multiple SNP haplotypes (**Figure 10**). In the latter context, grouping haplotypes according to the methylation state they produce may serve as a collapsing strategy that overcomes the typical power limitations of haplotype-based genetic analyses due to high dimensionality resulting from haplotype diversity. We implemented our suggested flag strategy on a publicly available dataset and observed it to be useful in order to identify additional probes of interest to the phenotype, many of which would have been removed by applying a typical SNP annotation removal strategy.

This gap hunting approach does not identify all SNP-affected probes, since it relies on detection of discrete clusters of percent methylation. SNP-affected probes may simply not have enough methylation variability to detect multiple modes even when they exist, or genotype-specific distributions may be so overlapping that discrete clusters cannot be detected. For the purposes of EWAS, methylation at CpGs related to the first scenario is not likely to be of interest since effects sizes with outcome will also be difficult to detect at low-variability CpGs. Probes characterized by the latter scenario would in theory be of interest in EWAS, but approaches for reliable identification of tightly overlapping distributions (but with distinct clusters) are limited. For example probes of this nature, we applied Gaussian mixture models with BIC and dip statistics, but could not consistently identify them as multi-modal and could not estimate the proper number of clusters. These methods show similar (limited) performance on both the beta value and M-value scales; the advantage of using beta values in gap hunting, however, is that the ‘threshold’ argument can be more easily informed by biological intuition. These alternative approaches could potentially be used to classify probes as having 1 versus more than 1 cluster, but this would result in many false positives, likely including many probes unaffected by SNPs, as described by Daca-Roszak et al. This is an unnecessary price to pay, considering that probes of these sorts of distributions do not seem to be prevalent to such a large degree, at least in SEED (**Figure 8**). To partially capture such probes, one could titrate the ‘threshold’ or ‘outCutoff’ input arguments to gap hunting to allow more liberal classification of gap probes.

Our work also emphasizes the utility of visual inspection of methylation distributions when considering the potential influence of SNPs on probe signals. Methylated and unmethylated signal intensities can be inspected with consideration of expected SNP effects (as in **Figures 2 and 3**). Such expectations are clear for C, G, and SBE SNPs, but less so for SNPs in the remainder of the probe. Some reference annotations specifically emphasize probes with SNPs less than 10bp away from the C site [22], but previous studies, based on SNP annotations, have found little [13] to no [16] effect of SNPs in the probe body. Yet, our results using study-specific genotype and 450k data, rather than reference data, found that SNPs at least up 7-8 bp away from the C site, and potentially up to 20 bp away, affect subsequent signal (**Figure 6**). While we focused on the distance to the C site of a single probe SNP, the multiplicity of SNPs in a single probe, and the specific number of base pairs affected, may also be influential. The complexity of analyzing this question with paired genotype and 450k data increases with the number of probe SNPs, as one needs to consider each combination of genotypes that could be encountered at all SNPs and compare the resulting signal from each of these groups.

We also note gap signals that are not due to an underlying SNP or variant. It is possible that a SNP could be influencing at least a fraction of these probes, as annotation schemes are imperfect in their lack of study specificity and may not account for rare variants. We did observed that among the gap probes not containing a known SNP, there was a greater proportion of 2-group gap signals compared to the gap probes likely attributed to SNPs. This is consistent with the signal expected if it were driven by rare variants. Other explanations for non-SNP-affected gap probes could include cell type heterogeneity specific to a genomic region, ambiguously mapping probes, or probes that fail via detection p-value; we found that approximately 21% of probes could be attributed to these factors. An additional explanation for these gap signals could be exposure or outcome associations for specific regions. This should be the focus of future work. Notably, this is also the goal of EWAS, further arguing for a “flag and consider” approach rather than removal.

The MethylationEPIC BeadChip, the next iteration of the 450k which queries over 850,000 CpG sites, is now being utilized for EWAS. Given that the Type I and Type II probe designs are retained in this new array, the gap hunting approach and influences of SNPs we have described will still be of importance. As a new subset of probes that merit special consideration in EWA studies, gap signals can help advance the field by providing insight into methylation signals mediating genetic signal, both locally and in a broader context.

### Conclusions

We demonstrate a new method, called ‘gap hunting’ to identify clustered distributions of methylation signal measured by the Illumina 450k platform. We apply this method in a peripheral blood DNAm data set, find that the identified ‘gap signals’ are mostly attributed to underlying SNPs, and demonstrate how specific SNP scenarios can lead to gap signal behavior. We also describe several implications of gap signals in EWA studies and emphasize their ability to serve as surrogates for the genetic background of the interrogated CpG site. We argue for gap signals as a new, study-specific, subset of 450k probes that merit special consideration in the EWAS pipeline.

## Methods

All analyses were carried out using R version 3.2.2 and *minfi* version 1.16.

### Study sample

The Study to Explore Early Development (SEED) is a multi-site case-control study of 3-5-year-old children with autism spectrum disorder (ASD), conducted in localities within 6 U.S. states. Approximately 2,600 families participated in SEED phase I, and the children were classified into three groups according to neurodevelopmental outcomes, as previously described. [23]. SEED participants consist of the case group with ASD and 2 control groups without ASD: a general population group and a developmental disabilities group. DNA methylation was measured on 610 children enrolled in SEED phase I. Genome-wide genotyping data were available on a subset (n=590) of children on whom the DNA methylation was measured (see Genotype Measurement section).

### DNA methylation measurement and quality control

Genomic DNA (gDNA) was isolated from whole blood samples using the QIAsymphony midi kit (Qiagen). For each of 610 gDNA SEED samples, 500 ng of DNA was bisulfite treated using the 96-well EZ DNA methylation kit (Zymo Research). Samples were then processed on the 450K Array, randomized across and within plates to minimize potential confounding effects introduced by batch. Illumina .idat files generated from the array were processed using the *minfi* R package for 450k Array data [24]. Standard pre-processing and QC measurements were performed, including the removal of bad arrays, replicate samples, and sex-discrepant samples, defined as those in which the predicted sex based on the *minfi* function ‘getSex’ was discordant with self-reported sex. Cell composition estimates were obtained via the ‘estimateCellCounts’ function, also in the *minfi* package. Additional samples with outlying cell composition estimates were removed. We did not perform any probe-level QC on these data, such as detection p-value or ambiguously mapping probe filtering [12], because we were interested in quantifying the degree to which they contributed to gap signal behavior. Finally, quantile normalization was performed as implemented in the ‘preprocessQuantile’ function in *minfi*. These processing steps resulted in 607 samples for use in the downstream analysis. Beta values (methylated signal/(methylated + unmethylated signal) +100) were calculated and implemented in downstream analyses.

### Genotype measurement, imputation, and quality control

Of the 607 samples in our DNAm dataset, 590 had whole-genome genotyping data available, which was measured using the Illumina HumanOmni1-Quad BeadChip. Standard QC measures were applied: removing samples with less than 95% SNP call rate, sex discrepancies, relatedness (Pi-hat > 0.2), or excess hetero- or homozygosity. Markers with less than a 98.5% call rate, or are monomorphic were removed. Phasing was performed using SHAPEIT [25] followed by SNP imputation via the IMPUTE2 software [26], and all individuals in the 1000 Genomes Project as a reference sample. Genetic ancestry was determined using EigenStrat program [21] and eigenvectors were utilized in statistical analyses, as described in detail below. Given our interest in the role of SNPs in producing gap signals, we limited all of our analyses to the 590 samples that had both genotype and 450k data. We also limited our analysis to those SNPs with a minor allele frequency ≥ 0.5%, as this value corresponded to the same number of individuals that would be allowed in the smallest gap signal group according to the default input arguments (see ‘gap hunting’ algorithm section).

### Identification of gap signals

We identified gap signals using a function we developed, called *gaphunter()*, that can be implemented using the *minfi* package [24].

A matrix of beta values (rows = probes, columns = samples) is input to or calculated by the function. The method incorporates two user-defined arguments: ‘threshold’, and ‘outCutoff’. For each row in this matrix:

1. Order beta values sequentially
2. Calculate consecutive pairwise differences for these values.
3. Determine the number of difference values calculated in 2) that are greater than the ‘threshold’ argument, defined herein as gaps. If one or more gaps exist, classify probe as gap signal and define the number of groups as the number of gaps plus one. If zero gaps exist, define probe as non-gap signal.
4. For all gap signals, use location of gaps to classify individuals into distinct groups.
5. (Optional) For all probes defined as gap signals, sum the number of samples in all groups except that of the largest count. Define ‘outlier-driven gap signals’ as those in which this sum does not exceed the user-defined ‘outCutoff’ parameter, which is the proportion of the total sample size (default value = 1%). Remove these outlier-driven gap signals from output.

We report the number of gap signals detected at all possible combinations of a series of threshold (0.025, 0.05, 0.10, 0.2) and outCutoff (0.005, 0.01, 0.05, 0.1) values. We chose to complete all subsequent analyses with results from setting the threshold argument to 0.05 and the outCutoff argument to 0.01. We also elected to implement this method on the Beta value scale because this allows for the threshold argument to be informed by biological intuition. In the case where a user only has M-values available, a simple transformation to the Beta scale can be performed [27].

### dbSNP138 and repeat element annotation

We developed an annotation of all polymorphisms that mapped to probes on the 450k Array in order to have available flexible information on the site(s) to which a polymorphism mapped (CpG site, SBE site, etc.) and on the polymorphisms themselves (minor allele frequency, etc.). The Database for Single Nucleotide Polymorphisms version 138 (dbSNP138), was downloaded from the UCSC Genome Browser [28]. All classes of polymorphisms in dbSNP138 were incorporated downstream: “single” (SNPs), “mnp” (multi nucleotide polymorphism), “microsatellite”, “insertion”, “deletion” and “in-del”. The latter three categories were grouped together to form a single “in-del” group. A final class of polymorphisms, called ‘range’, was created to lump together remaining dbSNP138 descriptions (“unknown”, “named”, “mixed”, etc.). We also downloaded a list of repeat elements from the UCSC Genome Browser generated via RepeatMasker [28]. We filtered this list to only include short and long interspersed nuclear elements, long terminal repeat elements, and simple repeats (micro-satellites). The ‘findOverlaps’ function in the R package ‘GenomicRanges’ was used to map the location of all annotated polymorphisms or repeat elements to the C, G, SBE, and probe locations of all 450k Array probes [29].

### Defining SNPs associated with C, G, and SBE sites

We were interested in analyzing the impact of specific SNPs (i.e. specific nucleotide changes) at specific locations in the probe (C, G, and SBE sites) through joint analysis of our SEED genotype and 450k data. We again used the ‘findOverlaps’ function in the ‘GenomicRanges’ R package to find which of our measured SNPs overlapped to the C, G, SBE and probe length sites of all 450k probes. We performed these overlaps separately for all 4 probe locations, and then pooled the overlap results together. We then removed probes that had more than 1 type of SNP mapping to them. For example, if in our overlap results we found that a probe had a mapping C site SNP and a probe-length mapping SNP, that probe was not considered in these analyses.

Once we defined a ‘clean’ set of probes with respect to the location at which they overlapped a SNP, we grouped together probes of similar relevant characteristics. For the C and G site SNP analyses, we grouped probes based on if they had the same: nucleotide change (C/T SNP, for example), probe design (Type I or Type II), SNP mapping location (C site or G site), and strand on which the CpG of interest is designed to be interrogated. All probe level information was found via the Illumina 450k manifest. For the SBE site SNP analyses, we grouped probes based on all of these criteria as well as the reference nucleotide of the SNP. This step was necessary in order to more easily understand in which genotypes to expect a loss of signal. Our groupings were done for all of the scenarios delimited in **Figures 2 and 3**.

For each of these scenarios, we collected 4 metrics across all probes that fell into that scenario, grouping samples by their genotypes at each probe. The 4 metrics were: percent methylation, methylated signal, unmethylated signal, and a copy number metric. The methylated and unmethylated signals were derived from the *minfi* function ‘getMeth’ and ‘getUnmeth’, which we performed on an un-normalized (output of *minfi* function ‘preprocessRaw’) R object. The copy number metric was defined as:

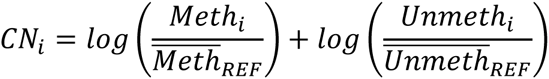

In this equation, ‘i’ refers to each individual and ‘Meth’ and ‘Unmeth’ to the methylated and unmethylated signals, respectively. At each probe the intensities are scaled by the mean values of the reference genotype of the SNP affecting that probe. This copy number metric therefore serves as way to jointly consider methylated and unmethylated signal, and more explicitly evaluate the difference between genotypes in terms of overall signal.

### Defining probe-associated SNPs

We were interested in evaluating the effect of distance from the C site to the SNP for situations in which a SNP was located in the probe but outside of the interrogated CpG and SBE positions. We first performed a similar overlap evaluation and filtering process as described above. Once we were limited to probes that had only overlapped measured probe-length mapping SNPs, we further filtered to probes that only had a single SNP in the probe length. This step was done in order to control for the potential effect of total amount of the probe length affected by SNPs. Next we grouped probes into bins of equal distance from the C site to the SNP, which was from 1 to 50 base pairs for the Type II design and 1 to 49 base pairs for the Type I design. At each probe, we identified the reference homozygote, heterozygote, and non-reference homozygote genotypes and group samples accordingly. We performed this grouping across probes within a specified distance value. Next we plotted the means and inter-quartile ranges (IQR; 25^th^ to 75^th^ percentiles) of the methylated and unmethylated signals as a function of distance, separately for the three genotype groups. The greater the discordance between the means and IQRs of the three groups indicated a greater effect of the mapping SNP.

### Defining probe categories

We were interested in comparing the overall standard deviation distributions of non-gap and gap signals. Moreover, we were further interested in within group differences relating to probe having an underlying SNP (measured or annotated) or not. For both non-gap and gap signals, we first identified probes that had at least one measured SNP anywhere in the probe body (through the same overlap analysis described above). From the remaining probes in each group, we identified which probes that overlapped at least one polymorphism in the dbSNP138 annotation described above, or a repeat element as defined by UCSC [28]. Again, overlap analysis in this case was also undertaken as described above. The remaining probes in each group were classified as having no underlying variant. This classification resulted in all 473,864 autosomal probes into 6 mutually exclusive categories: non-gap signals with no underlying (measured or annotated) SNP, non-gap signals with an annotated variant or repeat element, non-gaps signals with a measured SNP, gap signals with no underlying (measured or annotated) SNP, gap signals with an annotated variant or repeat element, and gap signals with a measured SNP. The distinction between a measured an annotated SNP underlying a probe is that of a SNP that we have complete certainty of existing in the SEED study population (as we imposed a MAF threshold of 0.5% in our 590 samples) compared to the existence of a SNP with some probability that is a function of MAF.

### Investigating additional sources of gap-like behavior

We were also interested in quantifying the role of additional factors in producing gap-like behavior in the gap signals that did not have a measured/imputed SNP or a mapping annotated variant via dbSNP138 or a mapping repeat element. We determined the proportion of these probes that were known to be ambiguously mapping, were determined to fail via detection p-value, or were previously determined to be cell-type distinguishing probes in whole blood. To define ambiguously mapping probes we used a previously defined list [12]. We defined a probe as a technical failure if more than 10% of samples had a detection p-value of greater than 0.01, as determined via the *detectionP()* function in the minfi package [24]. We defined a probe as distinguishing cell type if it had a p-value < 1E-8 reported from a previous study investigating differential methylation according to blood cell types [18].

### Comparing gap hunting to other methods to identify multi-modal distributions

We sought to investigate whether other methods that specifically identify multimodal distributions could overcome gap hunting’s insensitivity to distributions that appeared multimodal but did not cluster into discrete groups, but still retain the ability to identify methylation distributions that did have discrete groups. One key complication to this question is the fact that the ‘true’ status of methylation distributions at every measured probe is not known, which hampers the ability to assess the classification properties of alternative methods. To overcome this problem, we constructed a subset of 10,000 probes in which half were classified as gap signals by gap hunting (and were thus positive controls) and the other half were non-gap signals, did not have a measured/imputed or annotated SNP, and whose standard deviation was in the lowest decile of standard deviations across all autosomal probes (and were thus negative controls). In this way we could maximize our understanding of the true status of the probes we were testing as having a clustered distribution or not. The first method we tested was a Gaussian mixture model implemented via the ‘Mclust’ function in the *mclust* R package [30]. We allowed the function to select the best number of clusters (choice of 1 to 6) for each of the 10,000 probes based on a Bayesian information criterion. The second method we tested was the dip test in which the null hypothesis is that the data come from a unimodal distribution [19]; we implemented this test using the ‘dip.test’ function in the *diptest* R package [31]. We recorded the dip test p-value for each of the 10,000 probes and calculated the area under a receiver operating characteristic curve using the ‘auc’ function in the *MESS* R package [32] and generating dip test classifications at various p-value thresholds against the ‘true’ status of the 10,000 probes. We performed the mixture model and dip test experiments on both beta values and M-values (logit transform of beta values) to examine if the performances of these methods were affected by the scale of the methylation values.

### Population stratification

Upon confirming that gap signals were largely due to underlying SNPs, we were interested in exploring their potential to correct for population stratification. We calculated principal components (PCs) from gap signals and compared them to eigenvectors derived from GWAS, via the EIGENSTRAT method [21], and PCs derived from probes annotated with 1000 Genomes SNPs as described by Barfield et al. [20]. In the Barfield method, we used the option to include probes that directly overlapped with SNPs at the C site.

### Identification of variably methylated probes

We were interested in exploring gap signals in the context of a typical step in the EWAS pipeline to filter out probes that are of low variability. We calculated the standard deviation of all 473, 864 autosomal probes and calculated the percentages of gap and non-gap signals in the remaining probe set after imposing various standard deviation filters. Our cutoffs were ranged from the 5^th^ to the 99^th^ percentile of standard deviation across all probes.

### Relating gap signals to underlying haplotypes

We sought to demonstrate the potential for gap signals to serve as a surrogate for the local genetic sequence, on a haplotype scale. We phased our genotype data using the SHAPEIT software [25]. After downloading a list of recombination hotspots from the 1000 Genomes Project combined panel, we defined the locations between them as linkage disequilibrium (LD) blocks [33] and defined haplotypes from all of the measured SNPs within these LD block regions.

### Implementing gap hunting in an EWAS pipeline

We sought to illustrate the utility of incorporating gap hunting into a typical EWAS pipeline. We downloaded 450k data from a previous study examining placenta methylation and newborn neurobehavioral outcomes in 335 samples [34] from the Gene Expression Omnibus (GSE 75248) [35]. We performed functional normalization [36], and then removed samples according the following criteria: if the predicted sex via the getSex() function in the minfi package did not match the self-reported sex (N = 4), and samples with a detection p-value greater than 0.01 in more than 1% of probes (N =7). We removed probes at which more than 10% of samples had a detection p-value greater than 0.01 (n = 1,959), and if they were previously identified as being ambiguously mapping (n = 29,233) [12]. The resulting data included data on 454,502 probes and 324 samples. We performed ComBat to adjust for a known batch variable [37], and performed surrogate variable analysis to remove additional confounding due to cell type heterogeneity [38], in the absence of a reference panel of sorted placenta cell types. We did not remove probes that map to SNPs identified via reference annotation, in order to apply gap hunting on all cleaned probes. Finally, we removed probes mapping to sex chromosomes (as this was done in the previous study), identified gap signals via *gaphunter()*, and used the *limma* R package [39] to perform single-site association analyses relating DNAm to infant arousal, adjusting for gender and birthweight. We then noted the number of suggestively significant (p-value < 1E-4) hits, the number of these flagged as gap signals, and the number that would have been removed via SNP annotation, using a dbSNP137 annotation included in the *minfi* package.

## List of Abbreviations

BIC: Bayesian information criterion
dbSNP138: Database for single nucleotide polymorphisms, version 138
DNAm: DNA methylation
EWAS: epigenome-wide association study
MAF: minor allele frequency
PCs: principal components
SBE: single base extension
SEED: Study to Explore Early Development
SNP: single nucleotide polymorphism

## Acknowledgements

We would like to thank Arni Runarsson and Rakel Trygvadottir for their lab expertise and array processing.

## Additional Files

**Additional File 1 – Figure S1: Number of gap signals detected in SEED at various combinations of the ‘threshold’ and ‘outCutoff’ arguments to *gaphunter()*.**

**Additional File 2 – Figure S2: Examples of non-gap and gap signals found in SEED at 5%**

**‘threshold’ argument and 1% ‘outCutoff’ argument. Panel A**: 5 probes not identified as gap signals. **Panel B:** 5 probes identified as gap signals with 2 clusters. **Panel C:** 5 probes identified as gap signals with 3 clusters.

**Additional File 3 – Table S1: Distribution of group counts for gap signals in SEED.**

Breakdown of number of groups or clusters in the 11,007 gap signals found in SEED samples.

**Additional File 4 – Table S2: Breakdown of all C/G and SBE site measured polymorphism**

**scenarios.** We isolated specifics scenarios in which the following conditions were met: a probe contained a measured SNP that mapped to the C, G, or SBE sites of a probe, and it also did not contain any other form of mapping SNP. This table contains a list of all SNP C, G and SBE site scenarios herein and their corresponding Figure #. Also included is the number of probes analyzed for each scenario, along with the count and proportion of those probes that were classified as gap signals. Most probes in SEED that overlapped with measured SNPs were not classified as gap signals (though ~80% of gap signals did overlap with SNPs, see **Additional File 7**).

**Additional File 5 – Figures S3-S25: All Remaining C and G site scenarios for Type II and Type I probes.** Each additional scenario of a C and G site-mapping SNP delimited in Figure 2 not including the scenario show in Figure 3. Each of these figures contains the same panels (A-D) as seen in Figure 3 All scenarios demonstrate the expected behavior shown in Figure 2.

**Additional File 6 – Figures S26-S31: All Remaining SBE site scenarios.** Each additional scenario of a SBE site-mapping SNP delimited in Figure 4 not including the scenario shown in Figure 5. Each of these figures contains 4 plots, showing every combination of CpG site interrogations on the forward and reverse strand as well as which nucleotide is the reference nucleotide.

**Additional File 7 – Figure S32: The effect of SNPs located in Type I probes outside of the CpG or SBE position on methylated signal and unmethylated signal.** We examined specific scenarios in which the following conditions were met: a probe contained a measured SNP in the 50bp probe length, it also did not contain a SNP mapping to the C, G and/or SBE sites, and it contained only a single SNP in the probe length. We found all probes that met this criteria and varying values of distance from the SNP to the measured C site (1-50 bp). At each distance value, we plotted the mean and inter-quartile range of the people who were homozygous for the reference allele (‘Major Homozygote’), heterozygous (‘Heterozygote’) or homozygous for the minor allele (‘Minor Homozygote’). The degree of overlap between these 3 lines and their respective IQRs therefore demonstrates the effect of a polymorphism on subsequent 450k signal; the lack of overlap is directly correlated to an increased influence of the polymorphism. For both methylated signal (**Panel A**) and unmethylated signal (**Panel B**), polymorphisms at closer distance to the C site drive discordance between the 3 genotype groups. The relationship is less clear than for Type II probes, most likely because there are fewer Type I probes generally (and further fewer in this specific scenario) and the Type I design assumes that CpG sites within the probe length match that the methylation state of the interrogated CpG site. This assumption would be violated given our inclusion criteria for this analysis if the polymorphisms in question here occur at the C site of CpG site within the 50 bp probe length.

**Additional File 8 – Figure S33: MAF distributions of measured SNPs vs annotated SNPs that map to 450k probes.** We calculated the minor allele frequency (MAF) of all measured SNPs that mapped to gap signals, and determined the MAF for all of the annotated SNPs that map to gap signals as seen in the dbSNP138 annotation. The greater amount of SNPs with high MAF (>0.1) in the annotated SNP group may account for the higher area under the curve at higher standard deviation values as seen in **Figure 8**.

**Additional File 9 – Table S3: Group distributions of 3 different classifications of gap signals.** We compared the group distribution for the three groups – mapping measured SNP, mapping annotated SNP, and no mapping SNP – of gap signals. The two groups with mapping SNPs had a very similar relative proportion of groups, while the group with no mapping SNPs was comparatively enriched for distributions with 2 clusters or groups. This result lends additional rationale to a different mechanism besides SNPs as leading the gap signal behavior.

**Additional File 10 – Table S4: Alternatives to gap hunting do not correctly identify polymorphism-affected clusters.** For the probes shown in **Figure 7** and the gap signal in **Figure 1**, we explored other ways of identifying clusters. Specifically we examined a Gaussian mixture model clustering algorithm that selects an optimal number of clusters based on the Bayesian information criterion, and the dip test for unimodality (alternative hypothesis is that distribution is multi-modal). We recorded the number of clusters selected by the mixture model algorithm and the dip test p-value.

**Additional File 11 – Figure S34: Filtering on variably methylated probes at various cutoffs in the context of gap signals**. We calculated the proportion of gap and non-gap signals at various percentile thresholds of standard deviation cutoff (1% to 99%) to define a variably methylated probe. Researchers who filter on variable methylation prior to association analysis should be cautioned to be increasingly aware of gap signals (and subsequently their implications on DNAm related to disease described herein) as the cutoff to define a variably methylated probe increases.

